# FACT activity and histone H3-K56 acetylation promote optimal establishment of chromatin architecture independent of ongoing transcription in *Saccharomyces cerevisiae*

**DOI:** 10.1101/383984

**Authors:** Laura McCullough, Trang H. Pham, Timothy J. Parnell, Mahesh B. Chandrasekharan, David J. Stillman, Tim Formosa

## Abstract

FACT is a histone chaperone that can destabilize or assemble nucleosomes. Acetylation of histone H3-K56 weakens a histone:DNA contact that is central to FACT activity, suggesting that this modification could affect FACT functions. We tested this by asking how mutations of H3-K56 and FACT affect nucleosome structure, chromatin integrity, and transcription output. Mimics of unacetylated or permanently acetylated H3-K56 had different effects on FACT *in vitro* and *in vivo* as expected, but H3-K56 and FACT mutations caused surprisingly similar changes in transcription of individual genes. Notably, neither the changes in transcript levels nor the effects on nucleosome occupancy resulting from mutations conformed to the model that FACT is needed to overcome nucleosomal barriers during transcription initiation or elongation. Instead, the results suggest that both FACT and H3-K56ac are involved in establishing chromatin architecture prior to transcription and restoring it afterwards. They contribute to a process that optimizes transcription frequency, especially at conditionally expressed genes, and restores chromatin integrity after transcription, especially at the +1 nucleosome to block antisense transcription, but FACT appears to be less involved than expected in directly promoting transcription.

## Introduction

FACT (<u>FA</u>cilitates <u>C</u>hromatin <u>T</u>ranscription/<u>T</u>ransactions) is a conserved, essential histone chaperone that can both destabilize and assemble nucleosomes (1-4). FACT is a heterodimer of Spt16 with either Pob3 (in yeast and fungi) or SSRP1 (in higher eukaryotes), with each subunit comprising multiple histone-binding modules connected by unstructured linkers (4-6). SSRP1 has an intrinsic DNA-binding domain not found in Pob3, but both versions of FACT require additional freestanding DNA binding activity to support full activity (7, 8). The HMGB family member Nhp6 provides this function in yeast cells and *in vitro* (7-9). FACT is proposed to use these multiple domains to bind sites on nucleosomes, sequentially exposing and binding to additional buried sites to produce a dramatic, reversible change in the structure called a reorganized nucleosome (2, 4, 6, 7, 10). The strength of the histone-DNA contact at the entry/exit point of the nucleosome has a key role in this model, as breaking this contact is the first step towards reorganization, and forming it is among the last steps in the reverse reaction leading to nucleosome assembly.

Lysine 56 of histone H3 interacts with the phosphodiester backbone of the DNA at the entry/exit point of nucleosomes and acetylation of this residue weakens this contact (11). H3-K56 is acetylated in nascent H3-H4 dimers, so newly formed nucleosomes carry this mark globally after S phase (12-16). Hst3 and Hst4 deacetylate H3-K56 during G2/M, but subsequent nucleosome deposition restores the acetylated form, so the presence of H3-K56ac in interphase reveals sites of nucleosome turnover (14, 15, 17). Constitutive turnover appears to be an important mechanism for maintaining appropriate nucleosome occupancy in promoters (18), but it is not known whether persistent H3-K56ac at these sites is solely a consequence of turnover or if it has a functional role in promoting it, or what role turnover has in maintaining appropriate chromatin structure within genes.

Nucleosomes stall the progression of RNA Pol II *in vitro* and FACT was initially named for its ability to partially reverse this block (3, 19). If FACT is needed to allow RNA Pol II to proceed through each nucleosome it encounters as proposed in this model, FACT defects should cause stalled or inefficient polymerase progression *in vivo.* Attempts to detect a change in the rate of RNA Pol II elongation in FACT mutants have had mixed results, with some observing indirect effects (20), but others reporting no changes (21-23).

Several studies have reported FACT occupancy across transcribed regions in proportion to the frequency of transcription, supporting a role during transcription (24-27). This role could involve assisting RNA Pol II progression, but the occupancy profiles can also be explained by a more passive model in which FACT localization is driven by exposure of its binding sites during any process that disrupts nucleosomes, including transcription (5, 6, 28). FACT is proposed to promote transcription by destabilizing nucleosomal barriers to RNA polymerase progression, but depleting FACT caused increased nucleosome turnover in frequently transcribed genes, suggesting that when it is present it stabilizes nucleosomes during transcription *in vivo* (29). Stabilization has also been observed *in vitro* (19, 30), and several lines of evidence show that FACT is needed to restore nucleosome occupancy after transcription (14, 20, 24, 31-35). FACT has been implicated in destabilizing specific nucleosomes, but this involved removing nucleosomes from promoters during their activation, prior to transcription (22, 36-38). FACT therefore clearly influences chromatin structure and localizes to sites of ongoing transcription, but it remains unknown whether it promotes RNA Pol II progression *in vivo.*

FACT is essential for viability in many organisms, which has been attributed to central roles in both transcription and replication (2). However, emerging evidence indicates that differentiated cells of higher organisms remain viable and transcriptionally active with low levels of FACT (39, 40). Fundamental questions therefore remain regarding how FACT contributes to physiological processes, and under what circumstances its functions are essential.

Here, we investigated how FACT activity is affected by H3-K56 modification *in vitro* and *in vivo,* and examined genome-wide effects on transcript levels and chromatin structure. We reasoned that changing the strength of the histone:DNA contact at the nucleosome entry/exit point should alter FACT’s ability to reorganize nucleosomes, revealing which physiological functions depend on this activity. If reorganization contributes to overcoming nucleosomal barriers that limit transcription initiation or elongation, lower FACT activity should cause decreased initiation frequencies and reduced elongation rates. This should lead to decreased transcript output, especially from highly transcribed genes that depend on frequent initiation and long genes that depend on efficient transcription elongation. We also reasoned that global changes in nucleosome occupancy and positioning in mutants would reveal how reorganization contributes to chromatin structure.

Our results show that FACT and H3-K56 modification have important roles in establishing and maintaining the chromatin architecture that provides optimal transcriptional output, but they do not support a direct role for FACT in overcoming barriers to transcription initiation or elongation. Conditionally expressed genes with a “closed” promoter architecture (18) were most dependent on FACT to maintain normal transcript levels, consistent with a model in which FACT activity is most important for effecting transitions between active and repressed chromatin states. Both FACT activity and the ability to deacetylate H3-K56 were important for preventing antisense transcripts that initiated within genes from proceeding “backwards” through the +1 nucleosome, suggesting a novel role for this feature. Together, the results suggest that FACT is needed to establish or modify chromatin structure, and to maintain it after it is perturbed by transcription or other processes, but is less important for enabling transcription.

## Results

### Effects of H3-K56Q on FACT activity *in vitro*

Reorganization of nucleosomes by FACT has been detected *in vitro* through the formation of slow-migrating FACT:nucleosome complexes observed by native gel electrophoresis and by increased sensitivity to restriction endonucleases (41). The Spt16-Pob3(Q308K) mutant heterodimer is somewhat hyperactive for reorganization of nucleosomes but displays inefficient dissociation of the FACT:nucleosome complexes upon addition of competitor DNA (42 and Fig 1). This release defect has been interpreted as failure of a nucleosome assembly quality control step (7, 42). We previously found that specific mutations in histone H4 can suppress phenotypes caused by *pob3-Q308K in vivo* and also promote efficient dissociation of FACT(Q308K):nucleosome complexes *in vitro* (42), demonstrating that this assay accurately reports on a physiologically relevant function of FACT. In contrast, (Spt16-11)-Pob3 heterodimers are defective in producing reorganization, but release from complexes normally (43 and Fig 1). Its defects are also suppressed both *in vivo* and *in vitro* by specific H2A-H2B mutations, again validating these assays for measuring relevant functions of FACT *in vivo.*

**Figure 1.**
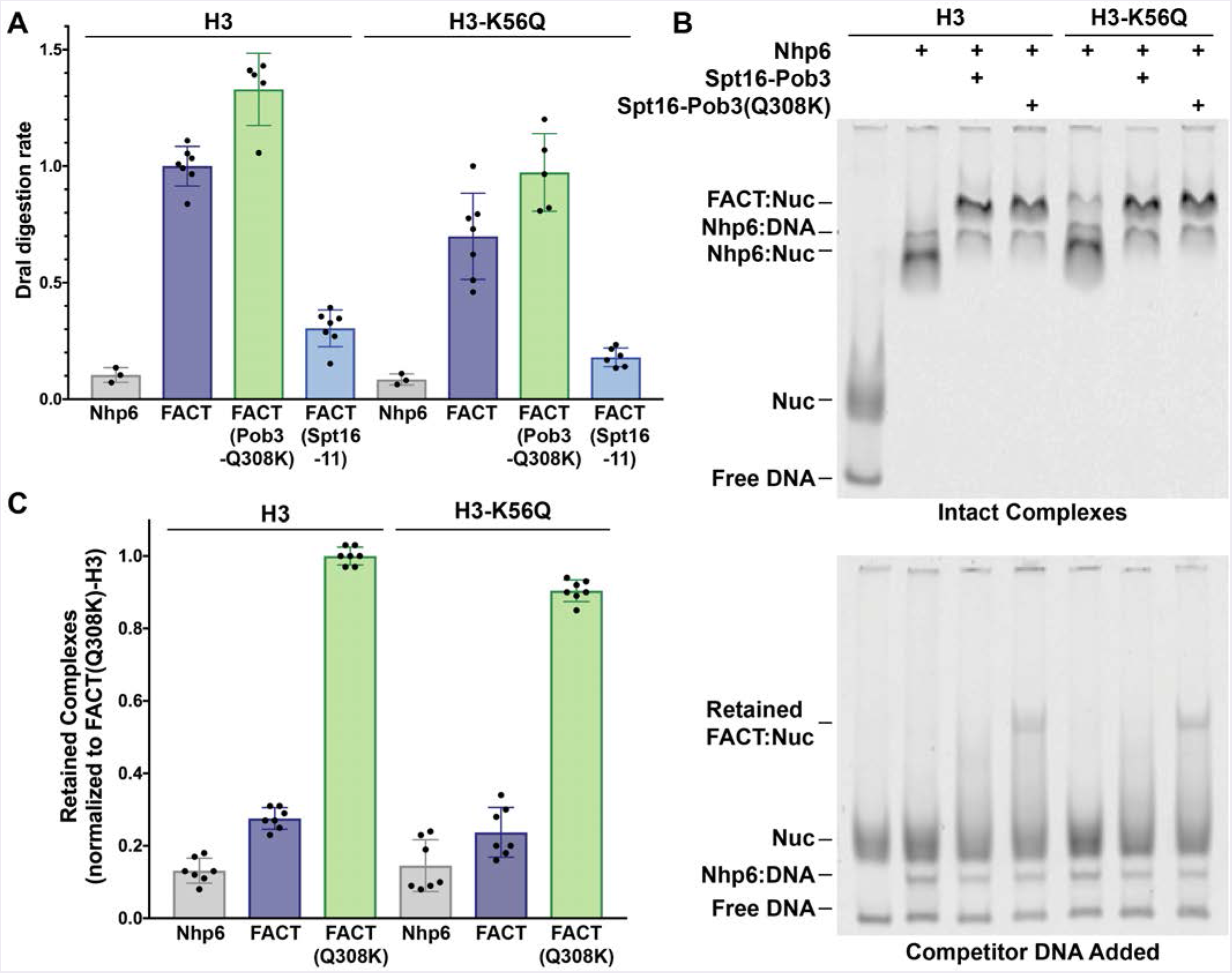
Effects of H3-K56Q on FACT activities *in vitro*. Nucleosomes were constructed with recombinant yeast histones (H3 or H3-K56Q) and a 181 bp 5S rDNA fragment as described previously (41). (A) The rate of DraI digestion at a site near the dyad was determined in the presence of 2 μM Nhp6 alone, or 2 μM Nhp6 with 0.2 μM Spt16-Pob3 (FACT), Spt16-Pob3(Q308K), or (Spt16-11)-Pob3. Initial rates of digestion were determined and normalized to the H3-containing nucleosomes treated with WT FACT. Error bars indicated the standard deviation. Comparing all conditions, H3-K56Q nucleosomes displayed an average rate of digestion that was 63% of the H3 nucleosome level and was different from WT with P < 0.0001. (B) Nucleosomes were tested alone (first lane) or mixed with Nhp6, Nhp6 + Spt16-Pob3, or Spt16-Pob3(Q308K) to form complexes (top panel), then unlabeled competitor DNA was added to initiate dissociation (bottom panel). (C) The fraction resistant to dissociation in (B) was calculated for 7 independent samples, with the average and standard deviation shown. Complex retention with Spt16-Pob3(Q308K) (labeled as FACT(Q308K)) with H3-K56Q was 90% of the level with H3 (P<0.0001).

To assess how weakening the histone:DNA contact at the entry/exit point affects reorganization by FACT, we assembled nucleosomes with H3-K56Q in place of H3. This has been validated as a mimic of H3-K56ac in a number of genetic and biophysical assays showing that it does destabilize the entry/exit point histone:DNA interaction (44-46). Nucleosomes constructed with recombinant yeast H3-K56Q bound FACT or its mutant versions normally but produced a lower rate of DraI digestion than normal, revealing a moderate defect as a substrate for reorganization (Fig 1A, 63% of WT averaged over all conditions). FACT(Q308K) produced the expected increase in digestion rate and deficiency in release from nucleosomes (7, 43), and this was weakly suppressed by H3-K56Q (Fig 1B). The suppression was statistically significant (Fig 1C) but minimal in magnitude (90% of the value with WT H3). We conclude that H3-K56Q drove the equilibrium away from the nuclease accessible form and towards nucleosome assembly, indicating reduced reorganization activity.

### Effects of H3-K56 status on FACT functions *in vivo*

To probe the correlation between the activities of FACT observed *in vitro* with its roles *in vivo,* we combined mutations that affect the status of H3-K56 with a range of FACT mutations. The commonly used *spt16-G132D* allele encodes an unstable protein that is rapidly degraded when cultures are shifted to a non-permissive temperature (24, 47). While useful for testing the effects of acute depletion of FACT, this allele is less informative for dissecting the importance of distinct functions of FACT in viable cells. We therefore tested the *pob3-Q308K* and *spt16-11* alleles that encode stable proteins with opposite effects *in vitro* (see Fig 1A) and distinct profiles of suppression *in vivo* (42, 43). We used three methods to alter the modification state of H3-K56: 1) loss of acetylation by deleting the machinery that writes this mark (the acetyltransferase *RTT109;* 12, 13, 48), 2) increased acetylation by removing the erasers of this mark (the deacetylases *HST3* and *HST4;* 14, 49, 50), and 3) mutating H3-K56 to alanine (uncharged, no interaction with DNA), arginine (constitutive stabilization of the DNA contact), or glutamine (permanent weakening of the contact). Histone H3 is produced in S. *cerevisiae* from two genes that encode identical proteins, *HHT1* and *HHT2* (51), so mutations are often tested by deleting the endogenous genes and supplying the mutated gene on a plasmid. This can result in copy number variation, and we have shown previously that this affects the phenotypes of FACT mutants and their interactions with histone mutations (42, 43). We also found that placing markers near the WT H3 genes caused slow growth when combined with FACT mutants (not shown), so we instead introduced the K56A, K56R, and K56Q mutations into both *HHT1* and *HHT2* through markerless conversion (52), adding translationally silent substitutions to allow detection of the alleles in genetic crosses (see Materials and Methods).

The H3-K56 mutations did not affect total H3 levels (Fig S1) and did not cause temperature sensitivity or sensitivity to hydroxyurea (HU) under the conditions tested (Fig 2; phenotypes consistent with those reported in 53 were observed under more stringent conditions; not shown). Combining *pob3-Q308K* with H3-K56 mutations caused synthetic defects ranging from severe with H3-K56A to minor with H3-K56R. The effects of H3-K56A or H3-K56R could reflect known overlapping functions of FACT with the histone chaperone Rtt106, which, unlike FACT, binds preferentially to (H3-H4)2 tetramers with the H3-K56ac modification (54-56). The severe growth defect caused by H3-K56A made genetic interactions difficult to interpret, so it was not studied further. H3-K56Q enhanced the growth defect, temperature sensitivity, and HU sensitivity caused by *pob3-Q308K,* while the effects of H3-K56R were more moderate and were limited to the HU sensitivity (Fig 2), suggesting a defect during DNA replication.

**Figure 2.**
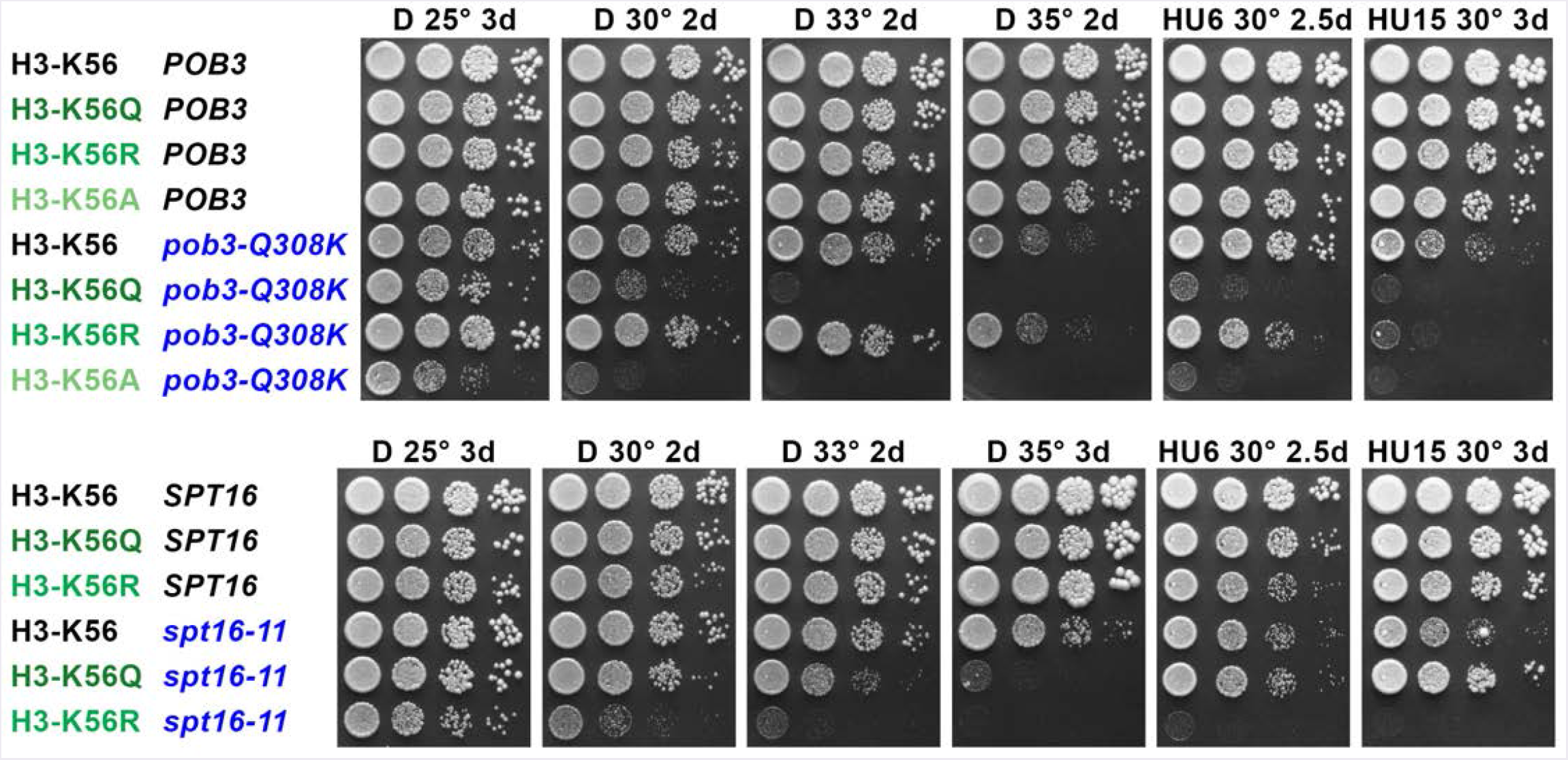
Genetic effects of combining H3-K56 variants with defects in different functional domains of FACT. H3-K56Q, K56R, and K56A alleles were constructed by markerless integration into the normal genomic *HHT1* and *HHT2* sites, then combined with alleles of FACT by standard genetic methods. 10-fold dilutions were tested for growth under various conditions as shown (see Materials and Methods). D is rich medium (YPAD). Labels indicate the concentration of hydroxyurea (HU, mM), the temperature, and the length of incubation in days. (Top panel) The temperature sensitivity (Ts^−^) and HU sensitivity (HUs) caused by *pob3-Q308K* were strongly enhanced by H3-K56Q or H3-K56A, while H3-K56R had no effect on the Ts^−^ phenotype and caused only moderate enhancement of the HUs. (Bottom panel) *spt16-11* displayed stronger synthetic defects with H3-K56R than with H3-K56Q at elevated temperatures, and a strong synthetic defect with H3-K56R on HU, but mild suppression of the HUs phenotype by H3-K56Q.

Combining *spt16-11* with H3-K56Q/R enhanced the Ts-phenotype, but in this case H3-K56R was more detrimental than H3-K56Q (Fig 2). Importantly, H3-K56Q moderately suppressed the HU sensitivity caused by *spt16-11.* These distinct genetic interactions are consistent with previous studies of these FACT alleles (42, 43), their opposite activity defects *in vitro* (Fig 1), and the expectation that H3-K56Q and H3-K56R would weaken and strengthen the entry/exit site contact, respectively. A weaker contact reduced reorganization activity *in vitro* (Fig 1) and was detrimental with FACT mutations, while a stronger contact partially suppressed a FACT allele with a defect in achieving reorganization. These results support the importance of reorganization as a core physiological function of FACT and demonstrate a role for H3-K56 status in this activity.

To compare the H3-K56Q acetylation mimic with the effects of H3-K56ac itself, we deleted the genes encoding the deacetylases Hst3 and Hst4 to produce increased levels of H3-K56ac (14, 49, 50, 57). This strongly enhanced the Ts- and HUs phenotypes caused by *pob3-Q308K,* just as the mimic H3-K56Q did (Fig S2A). Eliminating H3-K56ac by deleting Rtt109 (12, 13, 48) should mimic H3-K56R, and both manipulations had minimal effects when combined with *pob3-Q308K* (Figs 2, S2B). These results show that H3-K56ac affects the physiological functions of FACT, that being able to switch between stable and unstable configurations at this site is particularly important in FACT mutants, and that the reorganization may have a prominent role during DNA replication when nucleosome assembly rates are highest.

### FACT and H3 mutations caused similar defects in transcription

To determine how FACT and nucleosome reorganization participate in transcription, we used RNA-seq to measure the effects of chronic FACT defects and H3-K56 mutations on transcription. We then compared the expression changes for each mutant strain as log2 fold ratios relative to a wildtype strain (Fig 3). In addition to single FACT and H3-K56 mutations, we also tested a combined *pob3-Q308K* H3-K56Q strain (details listed in Table S1).

**Figure 3.**
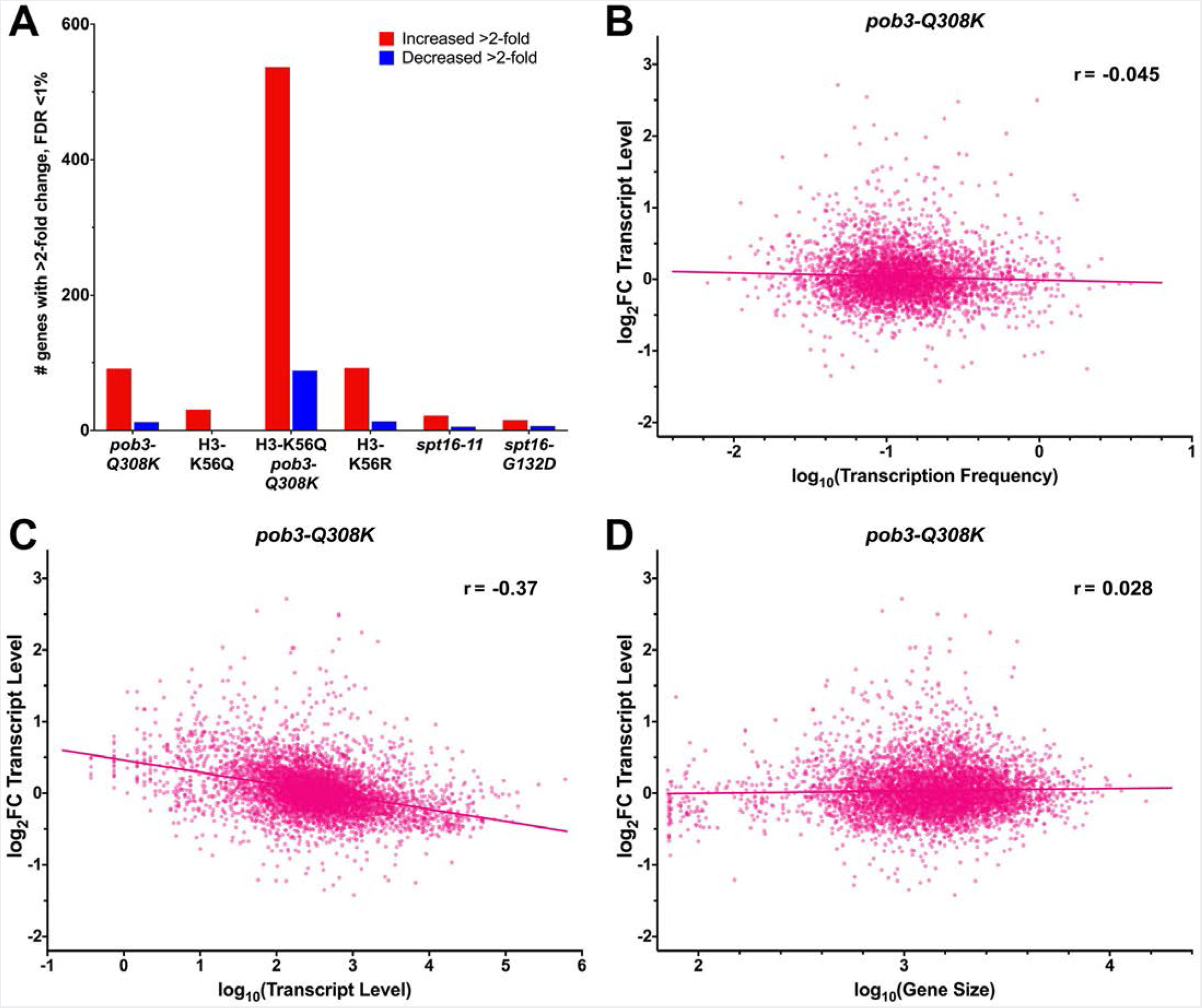
Global effects of mutations on transcript levels. Three biological replicates of WT and mutant strains were grown at 30° (strains used are listed in Table S1), and RNA-seq was performed as described in Materials and Methods. The log_2_(fold change) (log_2_FC) for each gene in the mutant relative to the WT was then calculated. (A) The number of genes with >2-fold changes in transcript level at a false discovery rate below 1% is shown for each strain. (B) The log_2_FC values for the *pob3-Q308K* FACT mutation are shown plotted against the log_10_ of the published transcription frequencies for each gene (58; see further explanation in the legend to Fig S3A). The Pearson correlation coefficient is given and a regression line plotted. (C) As in B, except the log_2_FC values are plotted against the log_10_ of the total number of transcripts observed in the WT strain or (D) against the log_10_ of the size of each gene in base pairs.

Each of the single mutants affected transcript output, and the double mutant had larger changes. Fig 3A shows the number of genes whose transcript level increased or decreased at least 2fold at a false discovery rate of 1%. The stability of the entry/exit point contact and some function of FACT therefore appear to have overlapping roles in transcriptional regulation. If that role involved promoting efficient initiation or elongation of transcripts, we anticipated that frequently transcribed genes would be differentially affected, as it would be difficult to maintain continued throughput at these sites if RNA Pol II were unable to maintain frequent initiation or continuous progress. However, neither FACT mutants nor H3 mutants displayed significant correlation between changes in transcript level and transcription frequency (Figs 3B, S3A). This analysis used a published dataset that combined RNA Pol II occupancy, transcript stability, and other factors to estimate the frequency of transcription for ~80% of yeast genes (26, 58). As a second test, we also compared changes in transcript level to the transcript abundance at steady state in our WT strain (Figs 3C, S3B, S3C). The latter method produced somewhat stronger statistical correlations (Pearson correlation coefficients from -0.20 to -0.37 for single mutants, -0.51 for H3-K56Q *pob3-Q308K*) but weak dose responses (a 10^6^-fold change in transcription frequency yielded about a 2-fold change in transcript level overall). Inefficient elongation should have a disproportionate effect on longer genes where efficient progression becomes more important, but this comparison also produced weak correlations and essentially flat dose responses (Fig 3D, S3D). These mutations therefore affected FACT’s reorganization activity and caused significant phenotypes, but they did not strongly affect the frequency of transcription initiation or the efficiency of elongation.

Pairwise comparisons of the changes in transcript level were used to ask how mutations affected individual genes (Fig 4; transcript level changes relative to WT for each mutant are shown compared to those of the *pob3-Q308K* strain). These comparisons produced strong correlations with positive, significantly nonzero slopes, revealing that expression of individual genes was affected similarly by different mutations. (Note that the *spt16-G132D* allele encodes an unstable protein whose functions are relatively normal under the permissive conditions tested here.) Combining *pob3-Q308K* with H3-K56Q produced effects that were roughly additive (Fig 4; compare panels A and C to panel B). This recapitulates the synthetic defects observed in genetic tests (Fig 2), and suggests genome-wide overlap of functions. We expected weakening and strengthening the entry/exit point contact to have opposite effects, but comparison of H3-K56Q and H3-K56R produced the highest correlation observed among pairs of mutants and a slope near 1 (Fig 4F). This suggests that H3-K56 acetylation itself does not directly affect transcript output, but the strength of this contact influences FACT activity globally.

**Figure 4.**
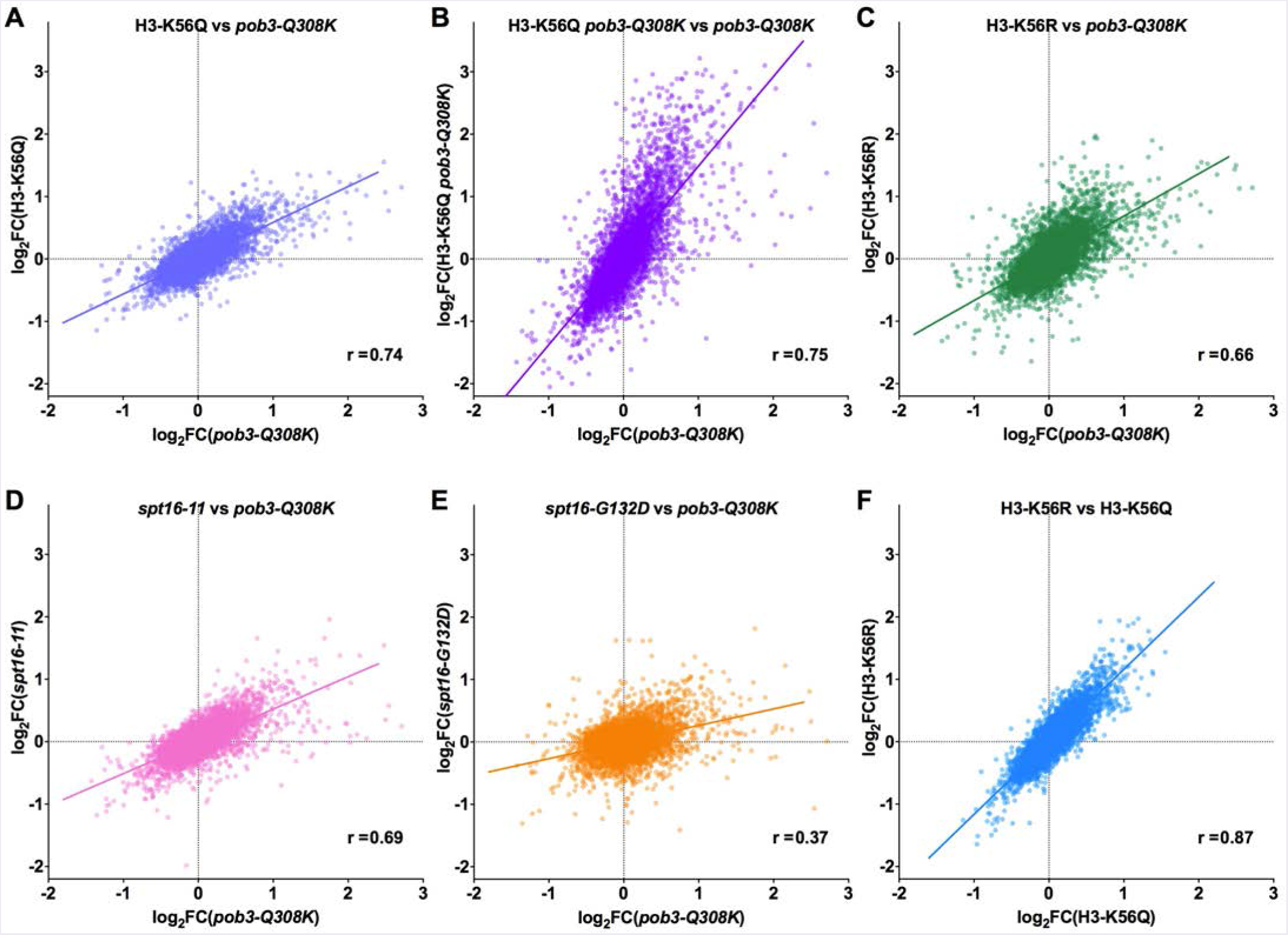
A range of mutations had similar effects on transcription of the same genes. The log_2_FC values for the change in transcript level relative to WT were calculated as in Fig 3, then plotted against each other in pairs. (A-E) Each mutant (y-axis) compared to the *pob3-Q308K* strain (x-axis). (F) The H3-K56R strain compared to H3-K56Q. The Pearson correlation coefficient is given and a regression line plotted.

### *pob3-Q308K* and H3-K56Q caused additive defects in the pattern of nucleosome occupancy

The similarity and additivity of the effects of FACT and H3-K56 mutations on transcript levels could reflect independent disruptions of a similar feature of chromatin structure. We performed MNase-seq with a subset of the strains to measure changes in nucleosome positioning.

To determine nucleosomal spacing around the transcription start site (TSS), we plotted the cumulative predicted midpoints of the nucleosomal fragments (calculated as 74 bp from the 5’ end of each sequenced fragment, assuming MNase digested all linkers efficiently). While the average change in positioning of the nucleosomes was negligible at the +1 nucleosome, a cumulative increase in spacing was observed downstream, primarily a 3’ shift of the nucleosome midpoint in *pob3-Q308K* mutants (Fig 5A). This effect was also observed in the H3-K56Q *pob3-Q308K* double mutant, along with decreased coherence of the peaks. Delayed redeposition of nucleosomes after passage of RNA polymerase might be predicted to shift nucleosomes downstream in the manner observed, with more frequent transcription leading to greater perturbation of the normal pattern. However, the expansion of nucleosome spacing was independent of the level of transcription, and so does not appear to be a response to RNA Pol II (Fig S5A, compare B to A and D to C). It has been estimated that the median gene in yeast is transcribed about once every eight minutes, and that fewer than 1 *%* of genes are engaged by more than one RNA Pol II molecule at the same time (58), making it less likely that transcription itself strongly affects nucleosome positioning. While the normal pattern of nucleosome positioning has mainly been attributed to inherent properties of DNA sequence and the effects of ATP-dependent remodeling (18, 59, 60), these results indicate that this histone chaperone also affects this property of chromatin.

**Figure 5.**
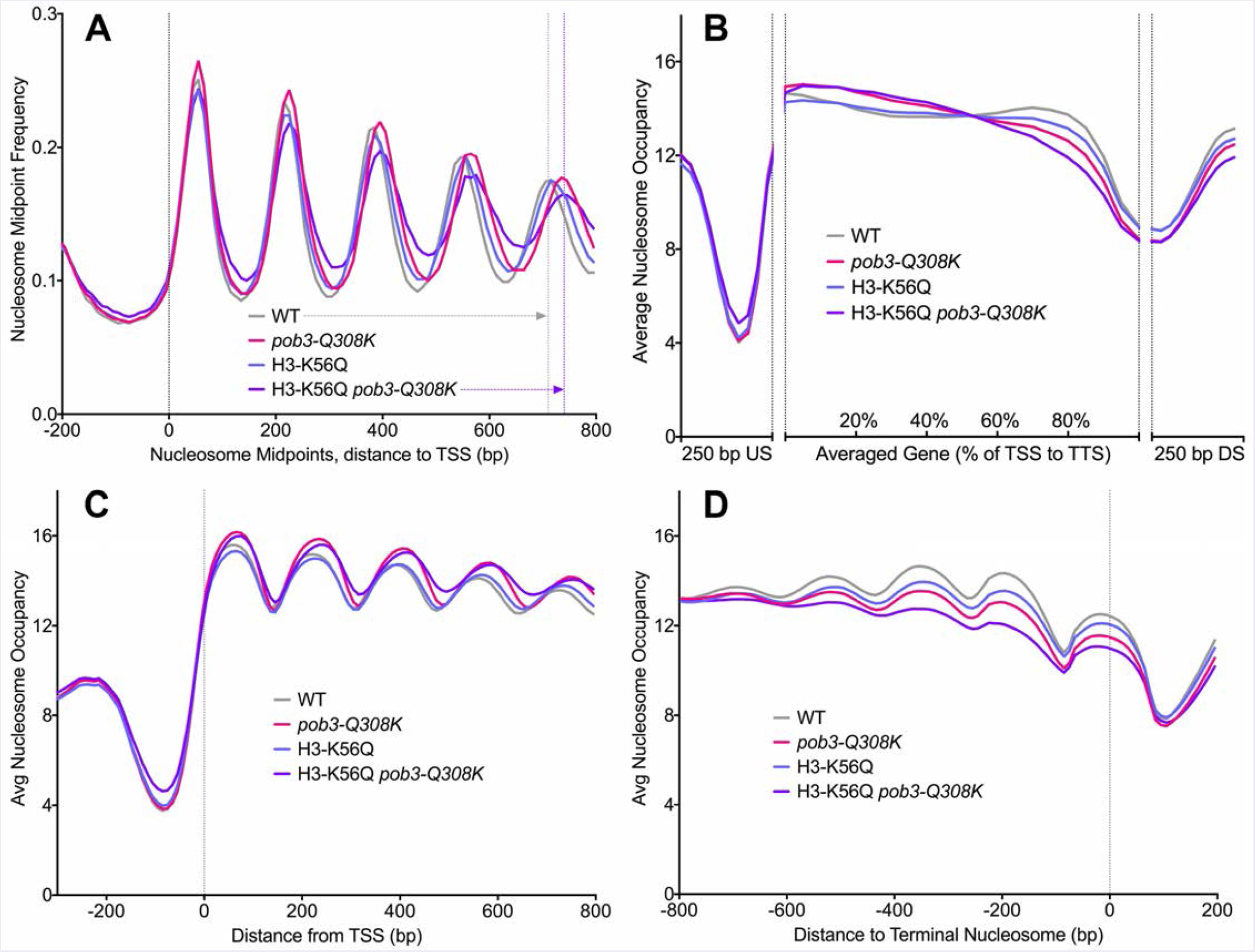
Effects of mutations on Nucleosome Positioning and Occupancy. MNase-seq was performed in triplicate for strains with the *pob3-Q308K* mutation, H3-K56Q, or the combination (see Table S1 for full genotypes). (A) Nucleosome midpoints were determined and aligned by the transcription start site (TSS) for each of 5337 genes. Vertical dashed lines indicate the accumulated displacement of 30 bp by the 4th nucleosome downstream of the TSS in the H3-K56Q *pob3-Q308K* strain relative to WT. (B) Average nucleosome occupancy was determined for each base. Each gene was normalized to the same size (100%) and the occupancy plotted in 40 bins. Occupancy is also shown for the 250 bp upstream (US) and downstream (DS) of each gene. (C-D) As in B, except genes were aligned by the TSS or by the last recognizable nucleosome in the transcription unit (the terminal nucleosome).

Defects in reorganization could cause either increased nucleosome occupancy (due to failed nucleosome destabilization) or decreased occupancy (due to failed tethering of nucleosomal components or lack of assembly activity). To test for these effects, we examined the distribution of nucleosome density (measured as the cumulative nucleosomal fragment depth where alignments were extended to 148 bp from each 5’ end) across a length-normalized gene body (Fig 5B). Occupancy increased at the 5’ ends of genes in *pob3-Q308K* mutants but decreased over the 3’ ends of genes in each single mutant, with additivity of the 3’ end effect in the *pob3-Q308K* H3-K56Q double mutant. While different models predict either increased or decreased nucleosome occupancy in the mutants, it is not clear how both effects would be observed in different regions of genes. For example, if a chronic defect in FACT activity causes increased probability of nucleosome displacement during each encounter of polymerase with a nucleosome, as inferred for acute FACT depletion (29), occupancy should decrease uniformly across individual transcription units in proportion to the frequency of transcription. However, the magnitude of the reduction of nucleosome occupancy over 3’ ends was independent of the level of transcription, so it does not appear to be a consequence of RNA Pol II passage (Fig S5B, panels A-F).

The decile of genes with the highest transcription frequency still displayed a skewed nucleosome distribution in mutant strains, but the pattern was different from that of average genes (Fig S5B, panels A-C). In the WT, highly transcribed genes had significantly lower than average nucleosome occupancy across the NDR, through the gene body, and into the downstream region. This remained true in the mutants but the effect was smaller, especially in gene bodies, meaning that the nucleosome occupancy in frequently transcribed genes increased as a result of the mutations. This could mean that frequent transcription displaces nucleosomes but having defective FACT/histones either 1) stabilizes nucleosomes against eviction, or 2) increases the efficiency of nucleosome deposition after eviction. Increased nucleosome turnover upon FACT depletion (29) suggests that option 1 is unlikely. H3-K56Q seemed to enhance nucleosome assembly by FACT, but Pob3-Q308K had the opposite effect (Fig 1A). In any case, enhanced deposition should be global, and should also have been observed in the regions flanking gene bodies. We speculate instead that highly transcribed genes are set up with a distinctive architecture that anticipates frequent transcription, including lower nucleosome occupancy over the gene body, and that FACT and H3-K56 mutations impair the ability to establish this pattern.

To remove potential distortion introduced by normalizing gene lengths, we aligned genes by their TSSs (Fig 5C), or by the nucleosome nearest to the transcription termination site (the terminal nucleosome; Fig 5D). These plots confirmed the 5’ increase/3’ decrease in nucleosome occupancy in mutants noted in Fig 5B, and showed that these effects extended to several nucleosomes near the TSS/TTS. The region immediately upstream of a yeast gene typically contains its promoter, and due to the dense packing of the yeast genome the region immediately downstream usually contains either the neighboring gene’s promoter or the transcription termination sites for both genes. We parsed the data by the orientation of the downstream neighbor and found that this did affect the patterns of nucleosome occupancy at the 3’ ends of genes (Fig S5C), but WT and mutants were similar, as were frequently and infrequently transcribed genes, so the orientation of the neighboring gene did not explain this effect. H3-K56ac and FACT therefore collaborate to establish or maintain nucleosome occupancy over the 3’ ends of genes, but neither their role in producing this feature nor its function were revealed by this analysis (see further discussion below).

### The relationship between altered nucleosome occupancy and changes in transcription

To examine how altered chromatin structure influenced transcription, we compared the change in nucleosome occupancy with the change in transcription of the gene. First, we examined nucleosome occupancy in the region from -30 bp to -150 bp upstream of the TSS to assess effects of changes just upstream of the initiation site (typically the gene’s promoter with low nucleosome occupancy). Second, we examined the region from the TSS to 130 bp downstream to determine the relevance of the +1 nucleosome. Third, we examined total occupancy over averaged gene bodies from the TSS to the TTS. We plotted nucleosome occupancy changes in each of these regions against the change in transcript level for each gene (Fig 6). Nucleosomes upstream of the TSS are expected to limit promoter function, and consistent with this occupancy of this region produced negative correlations with alterations in transcription (higher nucleosome occupancy was associated with decreased transcription; Fig 6, top row). However, the correlations and dose responses were moderate, with Pearson coefficients of -0.29 to -0.32 and a 2-fold change in transcript level being associated with about a 250-fold change in nucleosome occupancy.

**Figure 6.**
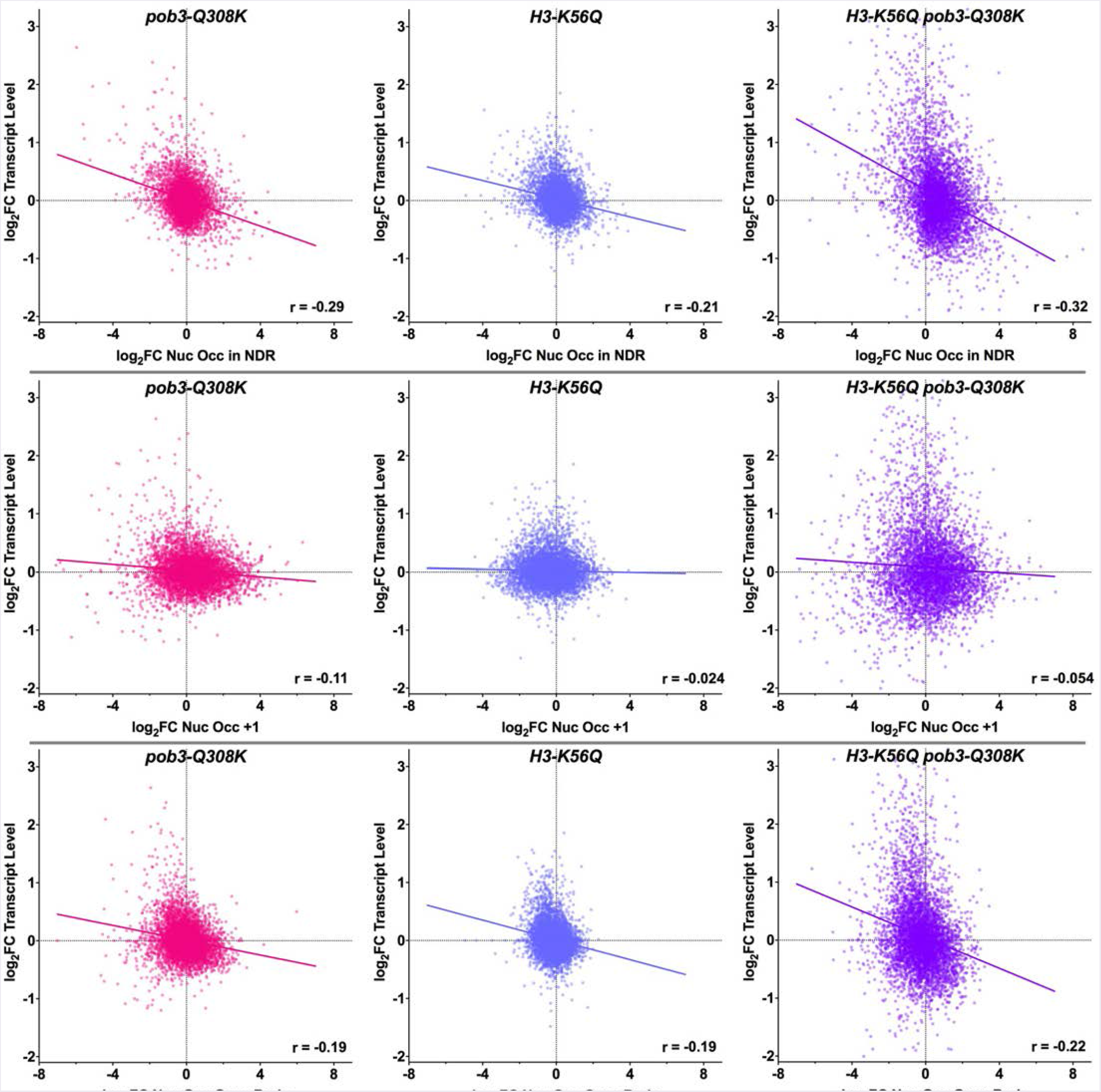
Changes in transcript level compared with changes in nucleosome occupancy in different regions of genes. The log_2_FC in nucleosome occupancy relative to the WT strain was determined for each mutant, then the average values were calculated for the nucleosome depleted region of each gene (NDR, 150 bp to 30 bp upstream of the TSS, top panels), the +1 nucleosome region (10 bp to 130 downstream of the TSS, middle panels), or the full gene body (bottom panels). These were plotted against the log_2_FC in transcript level for each gene, with the Pearson correlation coefficients and regression lines shown.

If FACT is required to allow RNA polymerase to overcome nucleosomal barriers, it would be expected to have a particularly important function at the +1 nucleosome as initiation transitions to elongation. Our results do not support this, as we observed very weak correlations between changes in nucleosome occupancy at this site and changes in transcription (Fig 6, middle row). Changes in nucleosome occupancy across the averaged gene body produced also produced low to moderate correlation with changes in transcript levels (Fig 6, bottom row). As noted above, the most highly transcribed decile of genes had lower nucleosome occupancy than average in WT cells (Fig S5B, panel A), but the global correlation was moderate when all genes were examined together (Fig S6A). Nucleosomes therefore were a barrier to transcription as expected, as decreased nucleosome occupancy in promoters was associated with a moderate increase in transcription, but changing the level of nucleosomes within genes, especially the +1 nucleosome, had less effect.

We next asked whether the genes experiencing the greatest increases in transcription displayed any similarities with one another. The top decile in the H3-K56Q *pob3-Q308K* strain (Fig S6B) had lower nucleosome positioning near the TSS (Fig S5A, panel E), higher than average nucleosome occupancy over what is normally the NDR (and also over the gene body and the downstream flank), and lower occupancy of the +1 nucleosome site (Figs S5B panel G, S6C). The lack of a prominent region of low nucleosome occupancy upstream of a subset of genes has been described as a “closed” promoter architecture, and is associated with conditional transcription, higher occupancy by both co-activating and co-repressing ATP-dependent remodelers, and high rates of nucleosome turnover in these promoters (18). These genes did not lose nucleosome occupancy over the NDR in the mutant strain, and decreased occupancy at the +1 nucleosome site was also observed in the decile of genes with the largest decrease in transcription (Fig S6C), so these features or changes in them do not appear to explain the changes in transcription. This shows that the repression produced by this distinctive chromatin architecture is particularly dependent on FACT activity and H3-K56 modification, but the changes that cause loss of repression of individual genes in the mutant strain are not visible after averaging large numbers of genes.

### H3-K56Q and H3-K56R have opposite effects on antisense transcription near the TSS

Depletion of FACT leads to increased aberrant transcription (24, 61). Our chronic FACT defects also caused increased abundance of antisense transcripts, particularly near the TSS (Fig 7). The antisense signal in both WT and mutant strains was slightly higher in genic regions than in promoter regions, with a distinct maximum near the average location of the +1 nucleosome, suggesting that this nucleosome may have caused stalled progression of this aberrant form of transcription. Again, combining *pob3-Q308K* with H3-K56Q caused a more severe phenotype. Notably, while H3-K56Q and H3-K56R caused similar levels of antisense transcripts upstream of the TSS, they had opposite effects near the +1 nucleosome, with H3-K56R producing low levels similar to the WT strain and H3-K56Q aligning with the more severe single mutants in FACT. This suggests that the +1 nucleosome has an antisense barrier function, that stabilizing the histone:DNA contact (permanently with H3-K56R or conditionally by deacetylating H3-K56ac) strengthens this barrier, and that FACT activity is needed to form this barrier. This is a novel function for the +1 nucleosome, but is consistent with its proposed roles in regulating sense transcription. Notably, in this context FACT enhances the barrier function of a specific nucleosome, the opposite of facilitating transcription. This is consistent with our overall conclusion that FACT is responsible for establishing optimal chromatin architecture, whether the goal is to promote or restrain transcription.

**Figure 7.**
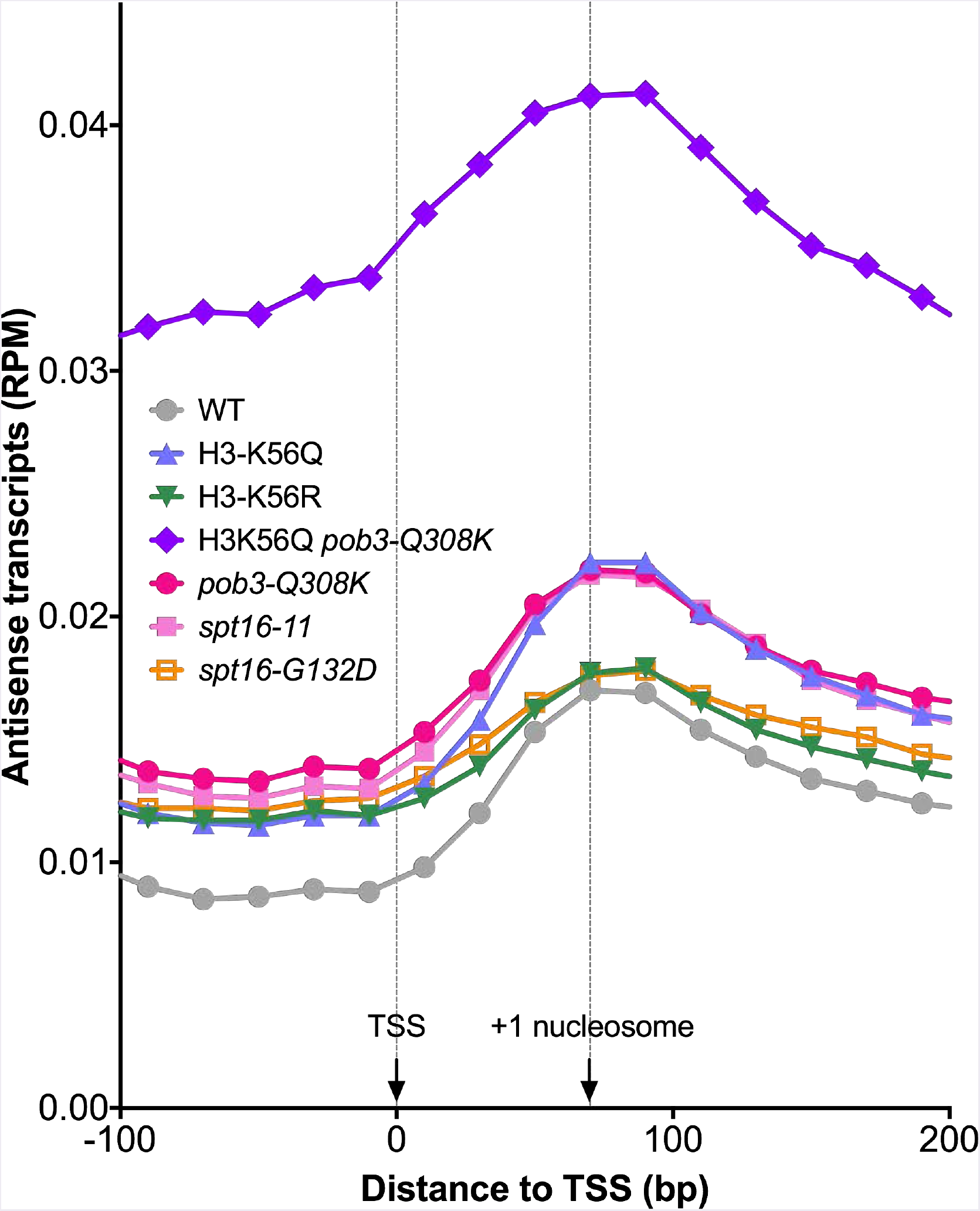
Effects of mutations on antisense transcription near the TSS. Transcripts were parsed into sense and antisense for ~5700 genes and the number of reads per million fragments (RPM) was calculated and mapped to a TSS-aligned metagene. The approximate location of the average +1 nucleosome is indicated at 70 bp downstream of the TSS.

## Discussion

The contact between histone H3 and DNA at the nucleosomal entry/exit site has a key role during the reorganization and assembly cycle promoted by FACT, so altering the strength of this interaction by manipulating H3-K56 acetylation status was expected to affect FACT’s activities *in vitro* and its functions *in vivo.* We found that a permanent mimic of H3-K56ac that weakens this contact made reorganization less favorable *in vitro,* tipping the reorganization equilibrium away from the open, nuclease accessible form and toward nucleosome assembly (Fig 1). This had opposite phenotypic effects on FACT alleles with different defects in promoting reorganization (Fig 2), confirming the importance of reversible control of H3-K56ac for optimal FACT function. Making reorganization more difficult is expected to be detrimental when combined with a mutation in FACT that already has a defect in promoting this transition, but a weaker entry/exit point contact could also complicate resolution of FACT:nucleosome complexes during nucleosome assembly, leading to either stalled or accelerated release, but causing decreased chromatin integrity in either case.

While different mutations had the expected opposing effects *in vitro* and *in vivo,* a broad range of mutations all had similar effects on transcription of individual genes (Fig 4). For example, H3-K56R and H3-K56Q should have opposite effects on the strength of the H3-K56:DNA contact, but they caused increased or decreased transcription of the same genes. Likewise, *pob3-Q308K* and *spt16-* 11, two alleles of FACT with distinct defects *in vitro* and different profiles of genetic interactions *in vivo* (42, 43) affected the same genes similarly genome-wide. The effects did not correlate well with the frequency of transcription or the length of the gene (Fig 3), so they do not suggest a defect in overcoming nucleosomal barriers during early stages of initiation or during elongation. Instead, we speculate that both FACT and H3-K56 acetylation contribute to a process that establishes chromatin architecture and restores it if it is disrupted. It is well established that nucleosome occupancy is lower in promoter regions and higher in genes; our results suggest that achieving precision in establishing these patterns and maintaining them requires FACT activity and that FACT activity is at least partly instructed or influenced by H3-K56 acetylation status. This process affects transcription and is affected by it, but is not limited to regions of ongoing transcription.

H3-K56Q and *pob3-Q308K* mutations also affected nucleosome positioning and occupancy profiles (Fig 5). Acute FACT depletion caused some loss of coherence of nucleosome positioning (24), and we also observed flattening of peaks and increased nucleosome spacing in genes (Fig 5A). More surprisingly, nucleosome occupancy increased over the 5’ ends of genes in a FACT mutant, decreased over 3’ ends in both FACT and H3-K56Q strains, and the 3’ end effects were additive (Figs 5C-D). Histone modifications are differentially distributed across transcription units, and histone and FACT mutations can cause increased FACT occupancy over the 3’ ends of some genes (62). Decreased nucleosome occupancy could therefore reflect an interaction between FACT and a histone modification, or between FACT and transcription termination. We can conclude that FACT activity and the properties of histones cooperate to establish or maintain nucleosome occupancy and positioning with different roles in different regions within genes. However, the changes in nucleosome occupancy had little correlation with either the normal level of transcription or the altered transcription in mutant strains, suggesting that they were not caused by transcription and were not regulating transcription frequency.

The correlation between FACT occupancy and the frequency of transcription has usually been interpreted as indicating that FACT is a component of the transcription machinery. The assumption, based on its activity *in vitro,* has been that it is recruited to promote passage of polymerases through nucleosomes. A different interpretation is that FACT binds to non-canonical nucleosomes, so its occupancy passively reflects sites where transcription, replication, repair, or the presence of modified or variant histones increase exposure of its binding sites. Consistent with this, FACT binds histone surfaces that are buried within canonical nucleosomes (5, 6), and FACT forms stable complexes with intermediates in nucleosome assembly or disassembly (28). In this view, localization is determined by the probability that FACT is converting an existing nucleosome to a non-canonical form, or it is bound to a non-canonical nucleosome to stabilize it or resolve it to assemble a nucleosome. The main drivers of FACT occupancy would then be signals to change the existing pattern of chromatin architecture to activate or repress conditionally expressed genes, and the requirement to restore chromatin integrity after perturbations caused by ongoing processes like transcription.

FACT defects caused the largest increases in transcription from a class of genes most dependent on chromatin structure, those with a “closed” architecture (18). This is consistent with a role in establishing global architecture. The inability to maintain repression by quickly restoring chromatin that has been disturbed by transcription is known to activate cryptic transcription (61) and FACT depletion leads to elevated antisense transcription (24). We also observed increased antisense transcription with FACT mutants, as well as with H3-K56 mutations (Fig 7). The distinct effects of H3-K56Q/R on antisense transcription match the expectation for opposite changes in nucleosome stability. The results suggest that deacetylating H3-K56 to strengthen the histone:DNA contact enhances the ability of the +1 nucleosome to block antisense transcription from proceeding into the gene’s promoter, and that normal FACT activity is needed to support this barrier function. In this case, FACT makes a nucleosomal barrier stronger, not weaker. Overall, our results support translating the acronym FACT as <u>FA</u>cilitates <u>C</u>hromatin <u>T</u>ransactions.

These results add to the accumulating evidence that FACT has a generic role in establishing and maintaining chromatin structure that is not limited to transcription. Data from mammalian cells suggests that this role may be more important during development than during proliferation of differentiated cells (39, 40), presumably because it is needed to support deconstruction of existing chromatin patterns and construction of new ones to allow transitions between transcription profiles.

## MATERIALS AND METHODS

Yeast strains are listed in Table S1 and are isogenic with the W303 background (63). Integrating *TRP1 orHIS3MX* markers downstream of WT *HHT1* and *HHT2* (42, 43, 64) caused slow growth of FACT mutants (not shown). We therefore used a method that integrated only the desired mutations in otherwise native genes (52). Translationally silent substitutions were introduced adjacent to the K56 target site, creating a SalI restriction site at codon 55, allowing the mutant allele to be detected by PCR amplification with primers specific to *HHT1* or *HHT2* (Table S2) followed by either endonuclease digestion or melting curve analysis (65). Accurate integration was confirmed by DNA sequencing, then standard genetic methods were used to construct strains with the desired combinations of mutations (66, 67). For phenotype tests, strains were grown to saturation in rich medium and 10-fold serial dilutions were plated on solid medium and incubated as described in each figure legend. MNase-seq and RNA-seq were performed in triplicate with the strains listed in Table S1, and bioinformatic analysis was performed as described (68, 69). Electrophoretic mobility shift assays (EMSAs), complex disruption with unlabeled competitor DNA, and restriction endonuclease sensitivity quantification were performed as described (41). Western immunoblots were probed with antibody to rabbit yeast histone H3 (Covance, special order using the peptide N-CKDIKLARRLRGER-C as the antigen) and mouse antibody to yeast Pgk1 (Molecular Probes), along with secondary antirabbit (800CW) and anti-mouse (680RD) secondary antibodies (Li-Cor), and scanned with an Odyssey CLx infrared scanner (Li-Cor).

## Acknowledgements

We thank Zaily Connell for assistance with western blots, and Emily Parnell and Robert Yarrington for contributions during the planning, execution, and writing of this manuscript. This work was supported by NIH grants R01 GM064649 to TF and DJS and R01 GM121079 to DJS.

Author contributions: LM and TP performed the experiments, TJP and TF performed bioinformatics analysis, TP and MBC prepared samples for genomic analysis, DJS and TF developed the experimental strategy and wrote the manuscript.

## Reference

1. Singer, R.A. and G.C. Johnston, The FACT chromatin modulator: genetic and structure/function relationships. Biochem Cell Biol, 2004. 82(4): p. 419–27. PMID: 15284894.

2. Formosa, T., The role of FACT in making and breaking nucleosomes. Biochim Biophys Acta, 2012. 1819(3-4): p. 247–55. PMID: 21807128;3229669.

3. Reinberg, D. and R.J. Sims, 3rd, de FACTo nucleosome dynamics. J Biol Chem, 2006. 281(33): p. 23297–301. PMID: 16766522.

4. Winkler, D.D. and K. Luger, The histone chaperone Fact: Structural insights and mechanisms for nucleosome reorganization. J Biol Chem, 2011. 286(21): p. 18369–74. PMID: 21454601.

5. Kemble, D.J., L.L. McCullough, F.G. Whitby, T. Formosa, and C.P. Hill, FACT Disrupts Nucleosome Structure by Binding H2A-H2B with Conserved Peptide Motifs. Mol Cell, 2015. 60(2): p. 294–306. PMID: 26455391; PMC4620744.

6. Tsunaka, Y., Y. Fujiwara, T. Oyama, S. Hirose, and K. Morikawa, Integrated molecular mechanism directing nucleosome reorganization by human FACT. Genes Dev, 2016. 30(6): p. 673–86. PMID: 26966247; PMC4803053.

7. McCullough, L.L., Z. Connell, H. Xin, V.M. Studitsky, A.V. Feofanov, M.E. Valieva, and T. Formosa, Functional roles of the DNA-binding HMGB domain in the histone chaperone FACT in nucleosome reorganization. J Biol Chem, 2018. 293(16): p. 6121–6133. PMID: 29514976; PMC5912460.

8. Valieva, M.E., G.A. Armeev, K.S. Kudryashova, N.S. Gerasimova, A.K. Shaytan, O.I. Kulaeva, L.L. McCullough, T. Formosa, P.G. Georgiev, M.P. Kirpichnikov, V.M. Studitsky, and A.V. Feofanov, Large-scale ATP-independent nucleosome unfolding by a histone chaperone. Nat Struct Mol Biol, 2016. 23(12): p. 1111–1116. PMID: 27820806.

9. Stillman, D.J., Nhp6: a small but powerful effector of chromatin structure in Saccharomyces cerevisiae. Biochim Biophys Acta, 2010. 1799(1-2): p. 175–80. PMID: 20123079; 2818483.

10. Hondele, M. and A.G. Ladurner, Catch me if you can: how the histone chaperone FACT capitalizes on nucleosome breathing. Nucleus, 2013. 4(6): p. 443–9. PMID: 24413069; PMC3925689.

11. Andrews, A.J., X. Chen, A. Zevin, L.A. Stargell, and K. Luger, The histone chaperone Nap1 promotes nucleosome assembly by eliminating non-nucleosomal histone DNA interactions. Mol Cell, 2010. 37(6): p. 834–42. PMID: 20347425.

12. Driscoll, R., A. Hudson, and S.P. Jackson, Yeast Rtt109 promotes genome stability by acetylating histone H3 on lysine 56. Science, 2007. 315(5812): p. 649–52. PMID: 17272722; 3334813.

13. Han, J., H. Zhou, B. Horazdovsky, K. Zhang, R.M. Xu, and Z. Zhang, Rtt109 acetylates histone H3 lysine 56 and functions in DNA replication. Science, 2007. 315(5812): p. 653–5. PMID: 17272723.

14. Kaplan, T., C.L. Liu, J.A. Erkmann, J. Holik, M. Grunstein, P.D. Kaufman, N. Friedman, and O.J. Rando, Cell cycle- and chaperone-mediated regulation of H3K56ac incorporation in yeast. PLoS Genet, 2008. 4(11): p. e1000270. PMID: 19023413; 2581598.

15. Kuang, Z., L. Cai, X. Zhang, H. Ji, B.P. Tu, and J.D. Boeke, High-temporal-resolution view of transcription and chromatin states across distinct metabolic states in budding yeast. Nat Struct Mol Biol, 2014. 21(10): p. 854–63. PMID: 25173176; PMC4190017.

16. Tsubota, T., C.E. Berndsen, J.A. Erkmann, C.L. Smith, L. Yang, M.A. Freitas, J.M. Denu, and P.D. Kaufman, Histone H3-K56 acetylation is catalyzed by histone chaperone-dependent complexes. Mol Cell, 2007. 25(5): p. 703–12. PMID: 17320445; 1853276.

17. Rufiange, A., P.E. Jacques, W. Bhat, F. Robert, and A. Nourani, Genome-wide replication-independent histone H3 exchange occurs predominantly at promoters and implicates H3 K56 acetylation and Asf1. Mol Cell, 2007. 27(3): p. 393–405. PMID: 17679090.

18. Parnell, T.J., A. Schlichter, B.G. Wilson, and B.R. Cairns, The chromatin remodelers RSC and ISW1 display functional and chromatin-based promoter antagonism. Elife, 2015. 4: p. e06073. PMID: 25821983; PMC4423118.

19. Hsieh, F.K., O.I. Kulaeva, S.S. Patel, P.N. Dyer, K. Luger, D. Reinberg, and V.M. Studitsky, Histone chaperone FACT action during transcription through chromatin by RNA polymerase II. Proc Natl Acad Sci U S A, 2013. PMID: 23610384.

20. Fleming, A.B., C.F. Kao, C. Hillyer, M. Pikaart, and M.A. Osley, H2B ubiquitylation plays a role in nucleosome dynamics during transcription elongation. Mol Cell, 2008. 31(1): p. 57–66. PMID: 18614047.

21. Mason, P.B. and K. Struhl, Distinction and relationship between elongation rate and processivity of RNA polymerase II in vivo. Mol Cell, 2005. 17(6): p. 831–40. PMID: 15780939.

22. Biswas, D., R. Dutta-Biswas, D. Mitra, Y. Shibata, B.D. Strahl, T. Formosa, and D.J. Stillman, Opposing roles for Set2 and yFACT in regulating TBP binding at promoters. EMBO J, 2006. 25(19): p. 4479–89. PMID: 16977311; 1589996.

23. Kuryan, B.G., J. Kim, N.N. Tran, S.R. Lombardo, S. Venkatesh, J.L. Workman, and M. Carey, Histone density is maintained during transcription mediated by the chromatin remodeler RSC and histone chaperone NAP1 in vitro. Proc Natl Acad Sci U S A, 2012. 109(6): p. 1931–6. PMID: 22308335; PMC3277555.

24. Feng, J., H. Gan, M.L. Eaton, H. Zhou, S. Li, J.A. Belsky, D.M. MacAlpine, Z. Zhang, and Q. Li, Noncoding Transcription Is a Driving Force for Nucleosome Instability in spt16 Mutant Cells. Mol Cell Biol, 2016. 36(13): p. 1856–67. PMID: 27141053; PMC4911744.

25. Mayer, A., M. Lidschreiber, M. Siebert, K. Leike, J. Soding, and P. Cramer, Uniform transitions of the general RNA polymerase II transcription complex. Nat Struct Mol Biol, 2010. 17(10): p. 1272–8. PMID: 20818391.

26. Pelechano, V., S. Jimeno-Gonzalez, A. Rodriguez-Gil, J. Garcia-Martinez, J.E. Perez-Ortin, and S. Chavez, Regulon-specific control of transcription elongation across the yeast genome. PLoS Genet, 2009. 5(8): p. e1000614. PMID: 19696888; PMC2721418.

27. Mason, P.B. and K. Struhl, The FACT complex travels with elongating RNA polymerase II and is important for the fidelity of transcriptional initiation in vivo. Mol Cell Biol, 2003. 23(22): p. 8323–33. PMID: 14585989.

28. Wang, T., Y. Liu, G.B. Edwards, D.D. Krzizike, H. Scherman, and K. Luger, The histone chaperone FACT modulates nucleosome structure by tethering its components. bioRxiv, 2018.

29. Jamai, A., A. Puglisi, and M. Strubin, Histone chaperone spt16 promotes redeposition of the original h3-h4 histones evicted by elongating RNA polymerase. Mol Cell, 2009. 35(3): p. 377-83. PMID: 19683500.

30. Chang, H.W., O.I. Kulaeva, A.K. Shaytan, M. Kibanov, K. Kuznedelov, K.V. Severinov, M.P. Kirpichnikov, D.J. Clark, and V.M. Studitsky, Analysis of the mechanism of nucleosome survival during transcription. Nucleic Acids Res, 2014. 42(3): p. 1619–27. PMID: 24234452; PMC3919589.

31. Biswas, D., R. Dutta-Biswas, and D.J. Stillman, Chd1 and yFACT act in opposition in regulating transcription. Mol Cell Biol, 2007. 27(18): p. 6279–87. PMID: 17620414; 2099615.

32. Voth, W.P., S. Takahata, J.L. Nishikawa, B.M. Metcalfe, A.M. Naar, and D.J. Stillman, A role for FACT in repopulation of nucleosomes at inducible genes. PLoS One, 2014. 9(1): p. e84092. PMID: 24392107; PMC3879260.

33. Ransom, M., S.K. Williams, M.L. Dechassa, C. Das, J. Linger, M. Adkins, C. Liu, B. Bartholomew, and J.K. Tyler, FACT and the proteasome promote promoter chromatin disassembly and transcriptional initiation. J Biol Chem, 2009. 284(35): p. 23461–71. PMID: 19574230.

34. Hainer, S.J., J.A. Pruneski, R.D. Mitchell, R.M. Monteverde, and J.A. Martens, Intergenic transcription causes repression by directing nucleosome assembly. Genes Dev, 2011. 25(1): p. 29–40. PMID: 21156811.

35. Hainer, S.J., B.A. Charsar, S.B. Cohen, and J.A. Martens, Identification of Mutant Versions of the Spt16 Histone Chaperone That Are Defective for Transcription-Coupled Nucleosome Occupancy in Saccharomyces cerevisiae. G3 (Bethesda), 2012. 2(5): p. 555–67. PMID: 22670226; PMC3362939.

36. Takahata, S., Y. Yu, and D.J. Stillman, The E2F functional analogue SBF recruits the Rpd3(L) HDAC, via Whi5 and Stb1, and the FACT chromatin reorganizer, to yeast G1 cyclin promoters. EMBO J, 2009. 28(21): p. 3378–89. PMID: 19745812; PMC2776103.

37. Takahata, S., Y. Yu, and D.J. Stillman, FACT and Asf1 Regulate Nucleosome Dynamics and Coactivator Binding at the HO Promoter. Mol Cell, 2009. 34(4): p. 405–415. PMID: 19481521; PMC2767235.

38. Shakya, A., C. Callister, A. Goren, N. Yosef, N. Garg, V. Khoddami, D. Nix, A. Regev, and D. Tantin, Pluripotency transcription factor Oct4 mediates stepwise nucleosome demethylation and depletion. Mol Cell Biol, 2015. 35(6): p. 1014–25. PMID: 25582194; PMC4333097.

39. Fleyshman, D., L. Prendergast, A. Safina, G. Paszkiewicz, M. Commane, K. Morgan, K. Attwood, and K. Gurova, Level of FACT defines the transcriptional landscape and aggressive phenotype of breast cancer cells. Oncotarget, 2017. 8(13): p. 20525–20542. PMID: 28423528; PMC5400524.

40. Garcia, H., D. Fleyshman, K. Kolesnikova, A. Safina, M. Commane, G. Paszkiewicz, A. Omelian, C. Morrison, and K. Gurova, Expression of FACT in mammalian tissues suggests its role in maintaining of undifferentiated state of cells. Oncotarget, 2011. 2(10): p. 783–96. PMID: 21998152; PMC3248156.

41. Xin, H., S. Takahata, M. Blanksma, L. McCullough, D.J. Stillman, and T. Formosa, yFACT induces global accessibility of nucleosomal DNA without H2A-H2B displacement. Mol Cell, 2009. 35(3): p. 365–76. PMID: 19683499; 2748400.

42. McCullough, L., B. Poe, Z. Connell, H. Xin, and T. Formosa, The FACT histone chaperone guides histone H4 into its nucleosomal conformation in Saccharomyces cerevisiae. Genetics, 2013. 195(1): p. 101–13. PMID: 23833181; 3761294.

43. McCullough, L., R. Rawlins, A. Olsen, H. Xin, D.J. Stillman, and T. Formosa, Insight into the mechanism of nucleosome reorganization from histone mutants that suppress defects in the FACT histone chaperone. Genetics, 2011. 188(4): p. 835–46. PMID: 21625001; 3176083.

44. Chen, C.C., J.J. Carson, J. Feser, B. Tamburini, S. Zabaronick, J. Linger, and J.K. Tyler, Acetylated lysine 56 on histone H3 drives chromatin assembly after repair and signals for the completion of repair. Cell, 2008. 134(2): p. 231–43. PMID: 18662539.

45. Watanabe, S., M. Resch, W. Lilyestrom, N. Clark, J.C. Hansen, C. Peterson, and K. Luger, Structural characterization of H3K56Q nucleosomes and nucleosomal arrays. Biochim Biophys Acta, 2010. 1799(5-6): p. 480–6. PMID: 20100606; PMC2885283.

46. Shimko, J.C., J.A. North, A.N. Bruns, M.G. Poirier, and J.J. Ottesen, Preparation of fully synthetic histone H3 reveals that acetyl-lysine 56 facilitates protein binding within nucleosomes. J Mol Biol, 2011. 408(2): p. 187–204. PMID: 21310161; PMC3815667.

47. VanDemark, A.P., H. Xin, L. McCullough, R. Rawlins, S. Bentley, A. Heroux, D.J. Stillman, C.P. Hill, and T. Formosa, Structural and functional analysis of the Spt16p N-terminal domain reveals overlapping roles of yFACT subunits. J Biol Chem, 2008. 283(8): p. 5058–68. PMID: 18089575.

48. Schneider, J., P. Bajwa, F.C. Johnson, S.R. Bhaumik, and A. Shilatifard, Rtt109 is required for proper H3K56 acetylation: a chromatin mark associated with the elongating RNA polymerase II. J Biol Chem, 2006. 281(49): p. 37270–4. PMID: 17046836.

49. Celic, I., H. Masumoto, W.P. Griffith, P. Meluh, R.J. Cotter, J.D. Boeke, and A. Verreault, The sirtuins hst3 and Hst4p preserve genome integrity by controlling histone h3 lysine 56 deacetylation. Curr Biol, 2006. 16(13): p. 1280–9. PMID: 16815704.

50. Maas, N.L., K.M. Miller, L.G. DeFazio, and D.P. Toczyski, Cell cycle and checkpoint regulation of histone H3 K56 acetylation by Hst3 and Hst4. Mol Cell, 2006. 23(1): p. 109–19. PMID: 16818235.

51. Smith, M.M. and O.S. Andresson, DNA sequences of yeast H3 and H4 histone genes from two non-allelic gene sets encode identical H3 and H4 proteins. J Mol Biol, 1983. 169(3): p. 663–90. PMID: 6355483.

52. Storici, F., L.K. Lewis, and M.A. Resnick, In vivo site-directed mutagenesis using oligonucleotides. Nat Biotechnol, 2001. 19(8): p. 773–6. PMID: 11479573.

53. Masumoto, H., D. Hawke, R. Kobayashi, and A. Verreault, A role for cell-cycle-regulated histone H3 lysine 56 acetylation in the DNA damage response. Nature, 2005. 436(7048): p. 294–8. PMID: 16015338.

54. Su, D., Q. Hu, Q. Li, J.R. Thompson, G. Cui, A. Fazly, B.A. Davies, M.V. Botuyan, Z. Zhang, and G. Mer, Structural basis for recognition of H3K56-acetylated histone H3-H4 by the chaperone Rtt106. Nature, 2012. 483(7387): p. 104–7. PMID: 22307274; PMC3439842.

55. Zunder, R.M., A.J. Antczak, J.M. Berger, and J. Rine, Two surfaces on the histone chaperone Rtt106 mediate histone binding, replication, and silencing. Proc Natl Acad Sci U S A, 2012. 109(3): p. E144–53. PMID: 22198837; PMC3271894.

56. Kemble, D.J., F.G. Whitby, H. Robinson, L.L. McCullough, T. Formosa, and C.P. Hill, Structure of the Spt16 middle domain reveals functional features of the histone chaperone FACT. J Biol Chem, 2013. 288(15): p. 10188–94. PMID: 23417676; PMC3624403.

57. Thurtle-Schmidt, D.M., A.E. Dodson, and J. Rine, Histone Deacetylases with Antagonistic Roles in Saccharomyces cerevisiae Heterochromatin Formation. Genetics, 2016. 204(1): p. 177–90. PMID: 27489001; PMC5012384.

58. Pelechano, V., S. Chavez, and J.E. Perez-Ortin, A complete set of nascent transcription rates for yeast genes. PLoS One, 2010. 5(11): p. e15442. PMID: 21103382; PMC2982843.

59. Lai, W.K.M. and B.F. Pugh, Understanding nucleosome dynamics and their links to gene expression and DNA replication. Nat Rev Mol Cell Biol, 2017. 18(9): p. 548–562. PMID: 28537572; PMC5831138.

60. Hughes, A.L., Y. Jin, O.J. Rando, and K. Struhl, A Functional Evolutionary Approach to Identify Determinants of Nucleosome Positioning: A Unifying Model for Establishing the Genome-wide Pattern. Mol Cell, 2012. 48: p. 5–15. PMID: 22885008.

61. Kaplan, C.D., L. Laprade, and F. Winston, Transcription elongation factors repress transcription initiation from cryptic sites. Science, 2003. 301(5636): p. 1096–9. PMID: 12934008.

62. Duina, A.A., A. Rufiange, J. Bracey, J. Hall, A. Nourani, and F. Winston, Evidence that the localization of the elongation factor Spt16 across transcribed genes is dependent upon histone H3 integrity in Saccharomyces cerevisiae. Genetics, 2007. 177(1): p. 101–12. PMID:17603125.

63. Thomas, B.J. and R. Rothstein, Elevated recombination rates in transcriptionally active DNA. Cell, 1989. 56: p. 619–630.

64. Toulmay, A. and R. Schneiter, A two-step method for the introduction of single or multiple defined point mutations into the genome of Saccharomyces cerevisiae. Yeast, 2006. 23(11): p. 825–31. PMID: 16921548.

65. Wittwer, C.T., G.H. Reed, C.N. Gundry, J.G. Vandersteen, and R.J. Pryor, High-resolution genotyping by amplicon melting analysis using LCGreen. Clin Chem, 2003. 49(6 Pt 1): p. 853–60. PMID: 12765979.

66. Rothstein, R., Targeting, disruption, replacement, and allele rescue: integrative DNA transformation in yeast. Meth. Enzymol., 1991. 194: p. 281–302.

67. Sherman, F., Getting started with yeast. Meth. Enzymol., 1991. 194: p. 3–21.

68. Ramakrishnan, S., S. Pokhrel, S. Palani, C. Pflueger, T.J. Parnell, B.R. Cairns, S. Bhaskara, and M.B. Chandrasekharan, Counteracting H3K4 methylation modulators Set1 and Jhd2 co-regulate chromatin dynamics and gene transcription. Nat Commun, 2016. 7: p. 11949. PMID: 27325136; PMC4919544.

69. Sdano, M.A., J.M. Fulcher, S. Palani, M.B. Chandrasekharan, T.J. Parnell, F.G. Whitby, T. Formosa, and C.P. Hill, A novel SH2 recognition mechanism recruits Spt6 to the doubly phosphorylated RNA polymerase II linker at sites of transcription. Elife, 2017. 6. PMID: 28826505; PMC5599234.

